# Mindin is essential for cutaneous fibrogenesis in a new mouse model of systemic sclerosis

**DOI:** 10.1101/2022.01.26.477822

**Authors:** Isha Rana, Sunny Kataria, Tuan Lin Tan, Edries Yousaf Hajam, Deepak Kumar Kashyap, Dyuti Saha, Johan Ajnabi, Sayan Paul, Shashank Jayappa, Akhil SHP Ananthan, Pankaj Kumar, Rania F. Zaarour, Haarshaadri J, Rekha Samuel, Renu George, Debashish Danda, Paul Mazhuvanchary Jacob, Rakesh Dey, Perundurai S Dhandapany, You-Wen He, John Varga, Shyni Varghese, Colin Jamora

## Abstract

Fibrosis is a result of chronically activated fibroblasts leading to the overproduction of extracellular matrix (ECM), causing tissue hardening and loss of organ function. Systemic sclerosis (SSc) is a fibrotic skin disease marked by inflammation, autoimmunity and vasculopathy along with progressive fibrosis of the skin and internal organs. A major bottleneck in understanding the etiology of SSc has been the lack of a holistic animal model that can mimic the human SSc disease. We found that the transcription factor Snail is overexpressed in the epidermis of SSc patients and a transgenic mouse recapitulating this expression pattern is sufficient to induce hallmark clinical features of the human disease. Using this mouse model as a discovery platform, we have uncovered a critical role for the matricellular protein Mindin in fibrogenesis. Mindin is produced by Snail transgenic skin keratinocytes and aids fibrogenesis by inducing inflammatory cytokine and collagen production in resident dermal fibroblasts. Given the dispensability of Mindin in normal tissue physiology, targeting this protein holds promise as an effective therapy for fibrosis.

## INTRODUCTION

Tissue fibrosis is a pathological condition in which the diseased or damaged tissue is replaced by excessive extracellular matrix (ECM), leading to tissue hardening and loss of organ function. Secretion of ECM by activated fibroblasts (myofibroblasts) is a physiological response of the wound healing program. However, unlike canonical dermal wound healing, in the context of fibrosis, fibroblasts remain persistently activated, and is likely sustained through positive feedback loops between fibroblast activation, inflammation (1), tissue stiffness (2), and vascular damage (3). When critical organs such as lungs, heart, kidney or liver become affected, the consequences of such over-scarring become fatal. For this reason, fibrosis is a major contributor to many diseases and ∼ 40% of mortalities worldwide are correlated with some form of underlying fibrosis (4).

Even though it poses a significant public health and socioeconomic burden, fibrosis still remains an irreversible condition with very limited treatment options, and this problem is compounded by the scarcity of biomarkers for disease susceptibility (5). Despite strong evidence for the pro-fibrotic activities of pathways such as TGF-β and endothelin (among others), clinical trials utilizing inhibitors of these signaling cascades have not lived up to the potential established in preclinical studies (4), (6). One explanation for this discrepancy is the presence of redundant pathways that can eventually bypass the inhibition of a single node, given the multifactorial nature of fibrosis. Thus, there remains important gaps in the understanding of the molecular and cellular features that underlie this pathology. In this regard, preclinical animal models that recapitulate both the temporal and multifactorial nature of fibrosis is important to unravel the molecular underpinnings of this complex disease and illuminate potential new routes of therapeutic intervention.

An example of a fibrotic disorder with poor prognosis is scleroderma, a group of skin fibrotic disorders with different subtypes classified by the extent of tissue involvement. The subcategories of scleroderma are - Localised scleroderma (LoS) and Systemic Scleroderma or systemic sclerosis (SSc). LoS is limited to affecting only certain areas of the skin (7). On the other hand SSc is a progressive fibrotic disease with inflammatory, autoimmune and vascular defects (8). Fibrosis characteristically initiates in the skin of distal extremities. The skin manifestations are limited to these areas in the case of limited SSc, however large areas of skin are involved in case of diffused SSc. Unlike, LoS, which negatively impacts the quality of life, SSc can progress to involve critical organs such as the lungs, heart and kidney often with fatal consequences. Many animal models have been developed to study SSc, and they have led to useful insights about the disease and represent different pathological and molecular subsets found in SSc (7) (9). However, none of these models can be considered holistic in the sense that SSc encompasses fibrotic, inflammatory, autoimmune and vasculopathic components in the skin followed by progression of the disease to internal organs. Currently, most mouse models such as bleomycin treatment, TGF-β1, TSK2 and TSK1, uPAR mouse models lack at least one of these components (10). The only mouse model that recapitulates all of these four aspects of SSc is the Fli1+/- Klf5+/- double heterozygote mouse model (11)(10). The Fli1+/- Klf5+/- model provides an excellent example of how creating a double haploinsufficient mice, which mimics the epigenetic suppression of Fli1 and Klf5 observed in SSc derived fibroblasts, can spontaneously lead to development of all cardinal defects of SSc followed by progression of fibrosis to the lungs. However, one drawback of such a global haploinsufficiency model would be the difficulty in distinguishing systemic effects from local ones.

Nevertheless, it is also important to characterize if these molecular and histological cross-species parallels can lead to a similar symptomatic course of disease development in the mouse. There is lacuna for mouse models which can faithfully reproduce the symptomatic pathogenesis along with the histological and molecular similarities to the human disease. A mouse model which can manifest the earliest symptoms of the disease such as Raynaud’s like phenomenon and puffy fingers (both indicative of vascular defects of SSc), followed by chronic inflammation, systemic autoimmunity, progressive fibrosis in skin and involvement of other internal organs would be a valuable tool to increase our understanding of the etiology of SSc and illuminate new ways to treat this disease.

## RESULTS

### The Snail transgenic mouse recapitulates clinical features of scleroderma patients

In light of the upregulation of Snail in multiple fibrotic tissues such as pulmonary fibrosis (12), renal fibrosis (13), and liver cirrhosis (14), and many skin fibrotic conditions (15), we analyzed the expression of this transcription factor in the human fibrotic skin disease scleroderma (SSc). qPCR analysis of skin biopsies collected from SSc patients revealed an average 5-fold increase in Snail expression relative to non-SSc (nSSc) skin biopsies (Figure 1a). Furthermore, we observed that expression of the Snail protein is localized to the basal layer of the epidermis in SSc patients (Figure 1b) whereas it is undetectable in nSSc skin. We previously developed a transgenic mouse with Snail under the control of the keratin 14 (K14) promoter which targets the transgene to the basal layer of the epidermis (16). This transgenic mouse recapitulates the expression pattern of Snail observed in human SSc skin (Figure 1b) and also develops dermal fibrosis (2), (17). Interestingly, although the RNA of the Snail transgene is expressed from neonatal to adulthood (Figure 1c), the protein is expressed only at the neonatal age (Figure 1d). This transient expression of the protein may reflect the inherent instability of Snail which often results in its rapid degradation (18) or changes in its subcellular localization (16). Consequently, a pulse of Snail expression in the epidermis is sufficient to induce dermal fibrosis. To test the extent that the overexpression of Snail in basal keratinocytes can induce the diagnostic characteristics of SSc, we compared the clinical manifestations observed in SSc patients with the phenotype of the Snail Tg mice skin. The Snail Tg mice were benchmarked against the clinical parameters as defined in the ACR-EULAR 2013 criteria (8).

**Figure 1:**
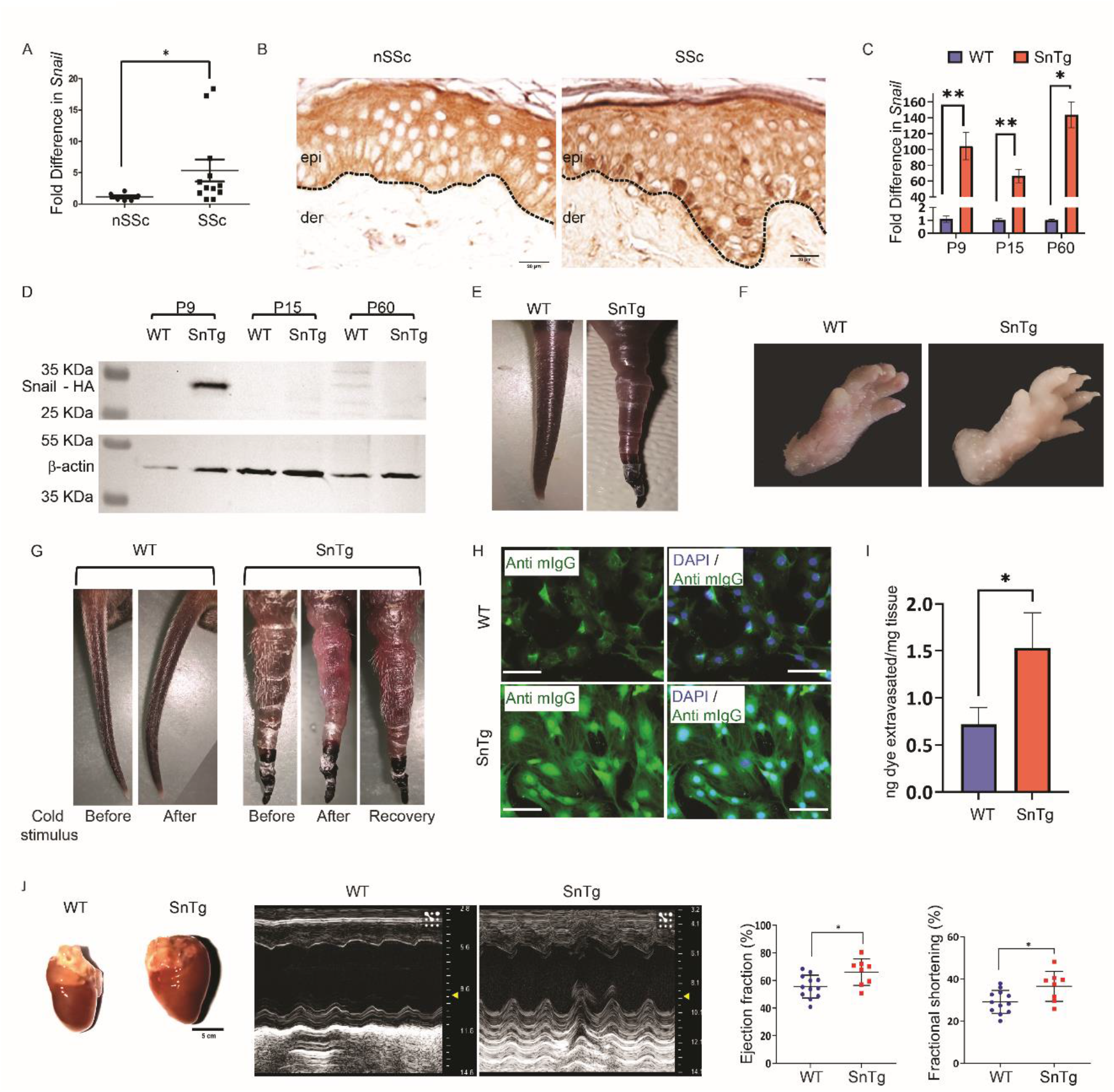
The Snail Tg mouse recapitulates diagnostic feature of scleroderma. A) qPCR for expression of Snail gene in biopsies taken from nSSc (n = 6) and SSc patients skin (n=12). B) Immunohistochemistry for Snail protein expression in skin sections taken from nSSc and from the skin of SSc patients. Black dotted line denotes the basement membrane that separates the epidermis (epi) from the dermis (der). C) qPCR for Snail expression in WT and Snail Tg mice skin samples taken at postnatal day 9 (P9) (WT n=4, Snail Tg n =6), P15 (WT n=5, Snail Tg n = 6) and P60 (WT n =6, Snail Tg n=6). D) Western blot for Snail using skin samples from P9, P15, and P60 WT and Snail Tg mice. Gross appearance of tail (E) and paws (F) of the WT and Snail Tg mice G) Cold challenge test for Raynaud’s phenomenon. Images of the tail for both WT and Snail Tg mice were taken before and after exposure to ice. Recovery was assessed in the Snail Tg animals 24 hrs post cold challenge. H) Anti-nuclear antibody staining using serum collected from P60 WT and Snail Tg mice (scale bar = 100 µm). I) Lungs of P60 WT and Snail Tg animals were analyzed for vascular leakage using dye extravasation assay (WT n =3, Snail Tg n=3). J) Comparison of gross anatomy (left-panel) and echocardiogram (right panel) of the heart of 1-year old WT and Snail Tg mice. Comparison of values for ejection fraction and fractional shortening obtained from echocardiogram between WT (n=12) and Snail Tg (n=8) mice (right panel). The error bars represent mean±SEM. P-value was calculated using Mann-Whitney’s U test (A), Welch’s t-test (C, I, J). (* p <0.05, ** p<0.01).

The most dramatic gross phenotypes in the neonatal Snail Tg mice (post-natal day 9), are puffy tails with necrotic ulcers, shiny skin (Figure 1e) and puffy paws with edema (Figure 1f). These features are similar to puffy fingers and necrotic lesions of digital ulcers observed in the tips of the fingers or toes in SSc patients (19). One of the earliest signs of SSc is Raynaud’s phenomenon, wherein the blood vessels in the extremities contract upon exposure to cold stress leading to the discoloration of the tissue (20). A brief cold challenge to the tail of neonatal mice (post-natal day 9) for 2 minutes, led to blanching of the tissue in the transgenic mouse that later recovered after 24 hours (Figure 1g).

One of the later hallmarks of the SSc that classifies the disease as an autoimmune disorder is the development of autoantibodies against nuclear proteins such as topoisomerase and the centromere (21). To test whether this autoimmune component observed in SSc patients is also present in Snail Tg mice, we tested for the presence of autoreactive antibodies in the serum by staining cells with serum collected from both wild type (WT) and Snail Tg mice. Whereas the WT serum from adult (P60) produced only background staining in primary dermal fibroblasts, the serum from the transgenic mouse contained antibodies against nuclear proteins (Figure 1h). This observation was not consistently reproducible in serum collected from postnatal day 9 mice. SSc is a progressive disease, in which fibrosis starts in the extremities but later involves internal organs. In line with this we observed that in adult (P60), the Snail Tg mouse revealed increased vascular leakage in the lung (Figure 1i), which is often an early sign of lung involvement in SSc (22). However, we did not observe any evidence of lung fibrosis at this age (Supplementary Figure S1a). Another major organ involved in scleroderma is the heart (23) and cardiac abnormalities usually portends poorly for the patient. Consistent with this clinical feature, echocardiography examination revealed that one-year old Snail Tg mice present with cardiac hypertrophy. This is manifested as increased heart size, increased left ventricle wall thickness, increased ejection fraction and fractional shortening, and decreased left ventricle volume at systole (Figure 1j and Supplementary Figure S1b). Together with the vascular leakage in the lung, the cardiac hypertrophy present in the year-old transgenic mouse may be secondary to the pulmonary defects that are observed in adult (P60) animals. Furthermore, in later stages of diffused SSc skin, skin ulcers are commonly observed over the extensor surfaces of the body (24). Consistent with this, we observed lesions in the dorsal neck region in ∼80% of year-old transgenic mice (Supplementary Figure S1c) that is often extended as the animal twists its head to groom itself.

In summary, these results highlight that five of the seven of the ACR-EULAR classification criteria for SSc were met in the Snail Tg mice (Supplementary table 1). The two criteria sclerodactyly and telangiectasia that were not reproduced is likely due to anatomical differences between the mouse and human. We did not observe nail fold capillaries in the transgenic animal, which may be due to the relatively thicker nails on the paws of the mice. In addition, sclerodactyly was not readily apparent as the mouse paws are naturally curled. Overall, the characterization of the Snail Tg mice demonstrates that transient expression of this transcription factor in the basal layer of epidermal keratinocytes is sufficient to launch the development of fibrosis. Moreover, similar to the clinical sequelae of SSc, the fibrotic phenotypes are long lasting in the transgenic mouse and eventually encompass the skin, connective tissue and internal organs.

### Snail Tg mice recapitulate histological and molecular characteristics of SSc

At the histological level, skin from SSc patients exhibits an increase in dermal thickness and loss of dermal white adipose tissues (dWAT) (25). While inflammation and dermal thickness may increase in the early inflammatory and edematous stages of the disease, the loss of dWAT is particularly associated with later, active fibrotic stages of SSc (25)(26). To test if the Snail Tg mice present with similar histopathological features, neonatal (P9) and adult (P60) skin from wild type(WT) and Snail Tg mice were analyzed. The dermal thickness of the Snail Tg skin was significantly increased by ∼1.5 fold relative to WT as early as P9, and increased to ∼3-fold at P60 (Figure 2a-b). However, the loss of dWAT was significant only in P60 transgenic skin (Figure 2a-b). Consistent with the increase in dermal thickness and loss of dWAT, we observed an increase in the amount of ECM proteins, collagen (Figure 2c) (2) and fibronectin (Supplementary Figure S2a), deposited in the dermis that is evident in the neonatal mice and substantially increases as the animal ages. The neonatal increase in ECM deposition is consistent with previous observations that activated fibroblasts are present in the dermis of the Snail Tg mouse as early as postnatal day 7 (17) along with increased expression of pro-fibrotic genes (2),(17). In sum, this data suggests that fibrogenesis initiates neonatally and progresses until it is most pronounced in the adult skin.

**Figure 2:**
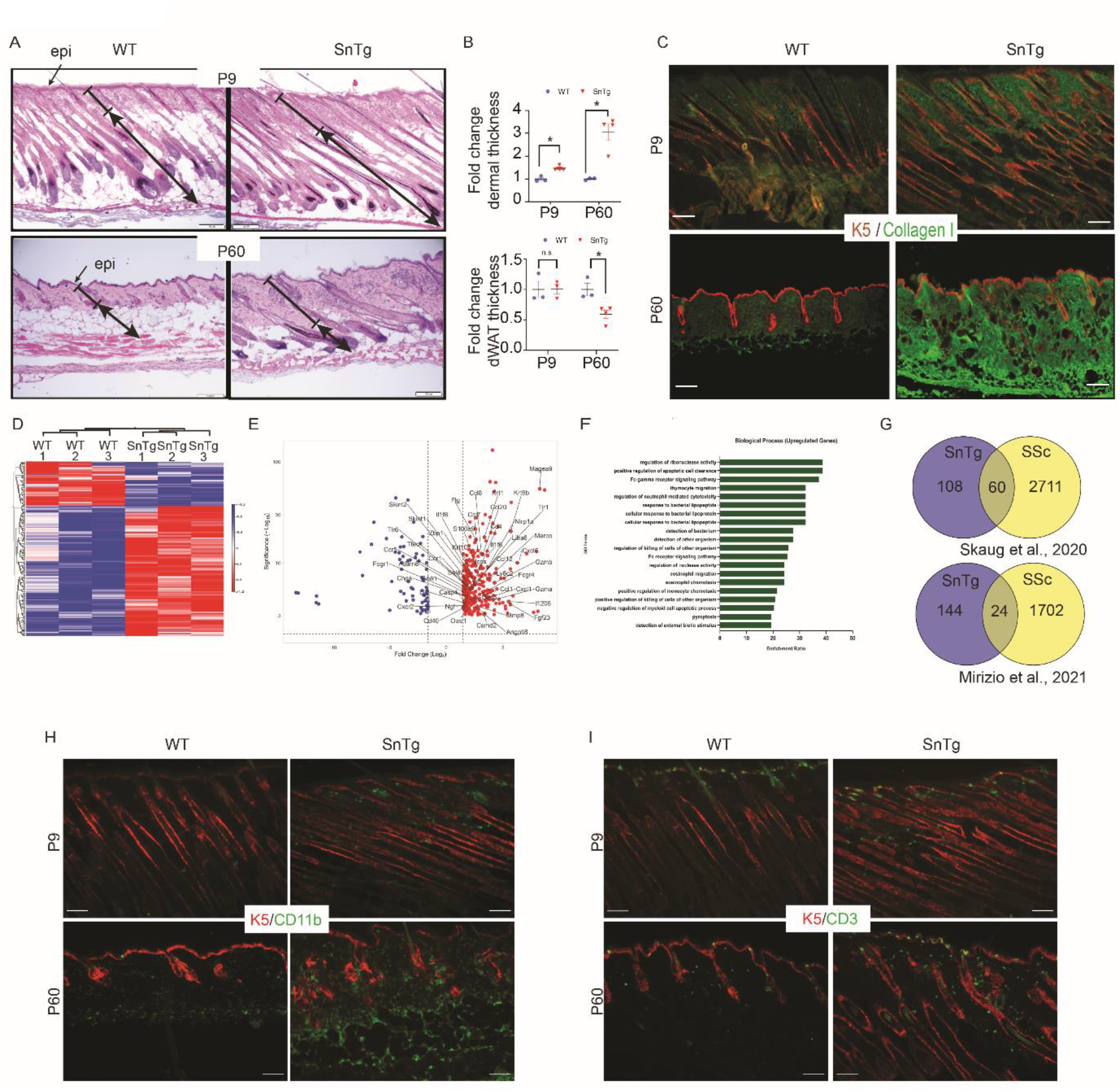
Snail Tg mice recapitulate histological and molecular characteristics of SSc. A) H & E staining of skin sections taken from WT and Snail Tg mice at P9 and P60 (scale bar = 100 µm). Block lines denote dermal thickness and lines with double headed arrows denote dermal white adipose tissue (dWAT) thickness B) Quantification of dermal and dWAT thickness in P9 WT (n=3) and Snail Tg (n=4), and P60 WT (n=3), Snail Tg (n=4) mice. C) Immunofluorescence imaging for Keratin-5 (K5, red) and Collagen 1 (green) in WT and Snail Tg mice at P9 and P60. D) Heatmap of differentially regulated genes in P9 WT (n=3) and Snail Tg (n=3) animals. Red = upregulated, Blue = downregulated E) Volcano plot of up and downregulated genes. F) Biological processes based on GO term enrichment of genes upregulated in the skin of the Snail Tg mice. G) Comparison between upregulated genes identified in RNAseq of P9 WT and Snail Tg mice with that of upregulated genes from SSc patients in Geo dataset[(GSE130955, (27)] and [(GSE166861, (28)]. H) Immunofluorescence imaging for K5 (red) and CD11b (green) in WT and Snail Tg mice at P9 and P60. I) Immunofluorescence imaging for K5 (red) and CD3 (green) in WT and Snail Tg mice at P9 and P60. The error bars represent mean±SEM. P-value was calculated using two-tailed Welch’s t-test (B). (* p <0.05).

Given the early onset of this phenotype, we investigated the process of cutaneous fibrogenesis in the Snail Tg mouse, by analyzing the transcriptome of P9 skin via RNAseq. A total of 353 genes were differentially expressed (DEG) (Log2 FC >1.5 for upregulated or <-1.5 for downregulated, Padj<0.05), and of these, 276 genes were upregulated and 77 were downregulated (Figure 2d, 2e). WEBGSALT analysis (27) revealed enrichment of GOTERMS associated with an inflammatory response (Figure 2f). This is consistent with earlier observations of a robust cutaneous inflammation in the Snail Tg mice at the neonatal level (2) (17). To test the extent to which Snail Tg mice skin is similar to SSc skin, we compared the skin transcriptome from P9 Snail Tg pups with the gene expression profile of SSc patients available in GEO datasets (28) (29). Of 168 upregulated genes in the Snail Tg mice with known human orthologs, 60 (35%) were also upregulated in early diffused SSc patients [GSE130955, (28)] (Figure 2g, 2% overlap, representation factor = 3.1, p<9.495e-17). Since the fibrosis in the Snail Tg mice begins at the neonatal age, we examined whether the gene profile of the Snail Tg mouse overlaps with pediatric localized SSc patients (29). We found 24 out of 168 (14%) orthologous genes upregulated in Snail Tg mice were also significantly upregulated in pediatric localized SSc patients’ skin [GSE166861(28)] (Figure 2g, 1.28%overlap, representation factor 2,p<7.724e-4)

Interestingly, the overlapping genes between the Snail Tg skin and human SSc skin were likewise enriched in inflammatory genes (Supplementary Figure 2b-2c). Consistent with the upregulation of inflammatory cytokine expression, we found a marked increase in immune cells recruitment upon immunohistochemical analysis of macrophages (Figure 2h), T cells (Figure 2i) and mast cells (Supplementary Figure 2d) in the Snail Tg skin. In support of increased infiltration of immune cells, upregulation of inflammatory cytokines and chemokines identified by the RNAseq analysis such as CXCL1, CCL12, CCL20, and S100A9, was verified by qPCR analysis of WT and Snail Tg skin at P9 and P60 (Supplementary Figure S2e-S2f). In addition, inflammatory signals found in SSc such as IL6, IL10, IL4, CXCL10, IL17F, and MCP1 (30),(31),(32),(33),(34),(35) were also upregulated in Snail Tg skin at P9 and P60 (Supplementary Figures S2e-S2f).

### Secreted Mindin from Snail transgenic keratinocytes mediates fibrogenesis

The central process in establishment of dermal fibrosis is the activation of fibroblasts (36). These fibroblasts undergo a phenotypic change from a normally quiescent state into a proliferative and contractile cell termed as myofibroblasts that are capable of secreting excessive amounts of extracellular matrix (ECM) proteins. In addition to depositing ECM, the myofibroblasts also secrete many different inflammatory cytokines. Given the varied activities of myofibroblasts, these cells have emerged as a coordinating node for the wide spectrum of processes that fuel fibrogenesis. Thus, the source of these activated fibroblasts has generated widespread interest in the hopes that interfering with their generation and/or maintenance would be an effective mode to prevent the onset and progression of fibrosis. Various competing hypotheses exist as to the biogenesis of these activated fibroblasts ranging from their differentiation from mesenchymal stem/progenitor cells (37) to epithelial-mesenchymal transition (EMT) (38) or an endothelial-mesenchymal transition (EndoMT) (39). The developmental role of Snail in mediating an epithelial-mesenchymal transition (EMT) during embryogenesis is widely extrapolated to be the mechanism by which activated fibroblasts are produced from neighboring Snail expressing epithelial cells. Surprisingly, using lineage tracing analysis, we found that none of the dermal fibroblasts in the transgenic skin were a product of an EMT (Figure 3a). The beta-Gal staining that we did observe in the dermis was co-localized with Oil-red O staining (Supplementary Figure S3a), indicating its localization in sebaceous glands owing to sebaceous gland hyperplasia that we have previously reported (40). This data suggests that Snail can activate resident dermal fibroblasts *via* the exchange of signals between the epidermis and dermis.

**Figure 3:**
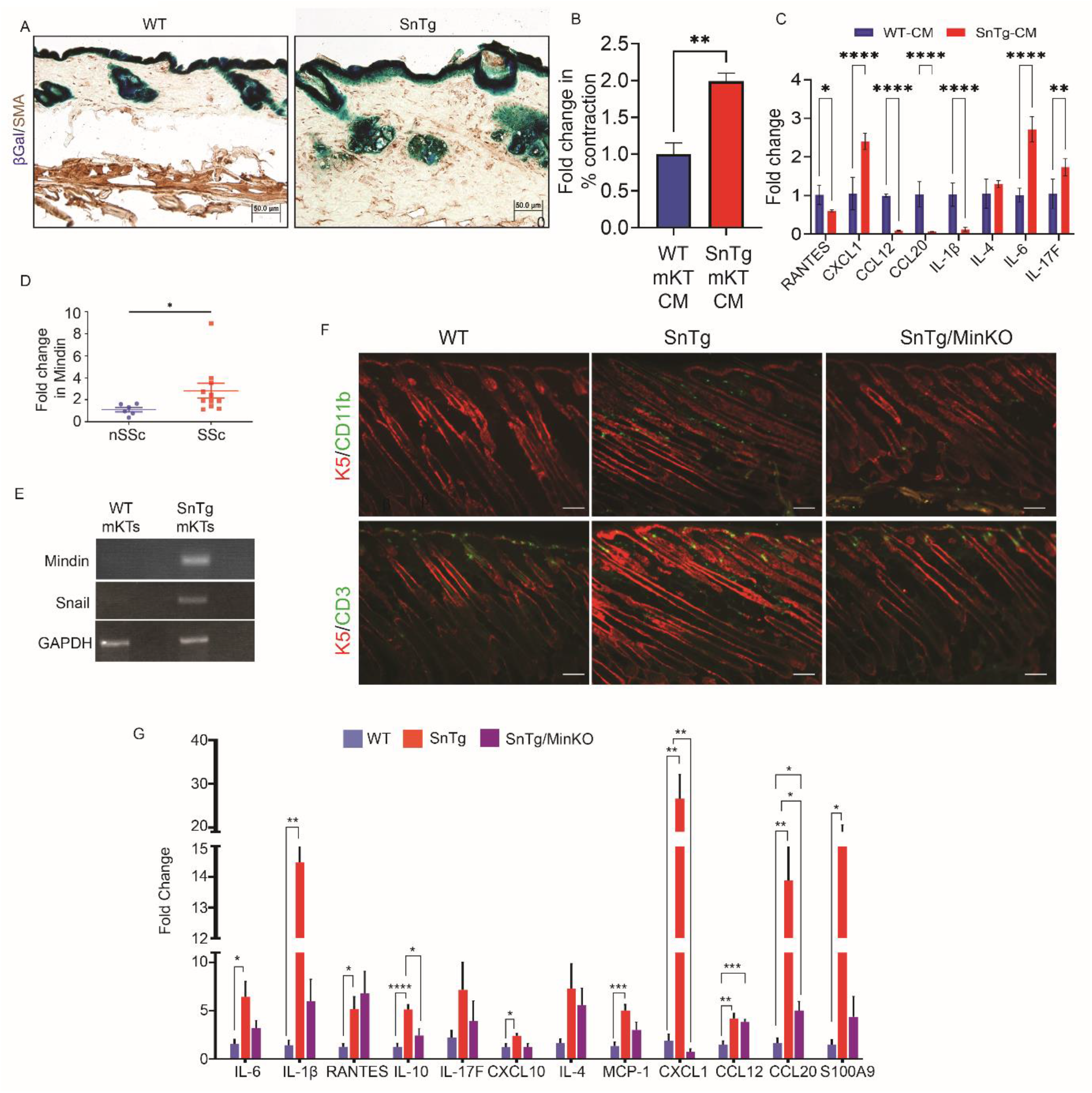
Secreted Mindin from Snail Tg keratinocytes, not an EMT, causes the early fibrotic phenotype. A. Lineage tracing of the WT and Snail Tg mice using the K14-Cre R26R reporter. All cells of epithelial origin are positive for β-Gal and stained blue (epidermis, hair follicles, and sebaceous glands) and activated fibroblasts are stained with α-smooth muscle actin (SMA, brown). B. Fibroblast activation measured with the collagen contraction assay. Conditioned media (CM) was collected from primary keratinocytes from WT (n=3) and Snail Tg (n=3) mice and added to a collagen plug embedded with primary dermal fibroblasts. C. qPCR for inflammatory cytokines gene expression in fibroblasts treated with WT (n=3) and Snail Tg (n=3) keratinocytes conditioned media. D. Expression of Mindin RNA in skin biopsies of nSSc (n=6) and SSc (n=12) patients. nSSc and SSc are the same patient samples used for Figure 1A. E. PCR for Mindin and Snail in primary keratinocytes from WT or Snail Tg skin. GAPDH is used as a loading control F. Immune cell recruitment in WT, Snail Tg, and Snail Tg/MinKO mice skin at age P9. Skin sections were stained for keratin-5 (K5, red) and either macrophages (CD11b, green) or T-cells (CD3, green). WT(P9) and Snail Tg(P9) are the same animals used earlier for Figure 2h-i G. qPCR for inflammatory cytokines gene expression in WT (n=8), Snail Tg (n≥6) and Snail Tg/MinKO (n=3) skin at P9. WT(P9) and Snail Tg(P9) are the same animals used earlier for Supplementary Figure 2e. The error bars represent mean±SEM. P-value was calculated using two-tailed Welch’s t-test (B,C), Mann-Whitney’s U test (D), and One way ANOVA followed Tukey’s post-hoc analysis for multiple group comparison (G) (* p <0.05, ** p <0.01, *** p <0.001, **** p <0.0001).

To test this hypothesis, we collected the conditioned media (CM) from WT and Snail Tg epidermal explants and assessed their sufficiency to activate primary dermal fibroblasts using a collagen contraction assay. Interestingly, Snail Tg epidermal conditioned media induced 1.7-fold higher activity relative to the WT CM (Supplementary Figure S3b). However, the epidermis is not a homogenous tissue of keratinocytes but also contains other cell populations including resident immune cells. To determine whether the transgenic keratinocytes are the source of the fibroblast activating signal(s), we used conditioned media from Snail Tg primary keratinocytes cultures and compared to media conditioned by wild type epidermal keratinocytes. We found that CM from Snail Tg cells was sufficient to induce collagen contraction two-fold compared to media conditioned from wild type keratinocytes (Figure 3b) and significantly upregulated markers of dermal fibroblast activation, αSMA (Supplementary Figure S3c). Furthermore, to test if CM from Snail Tg keratinocytes can also induce the inflammatory program in activated fibroblasts, we treated the primary dermal fibroblasts with either WT or Snail Tg CM and quantified the expression of inflammatory cytokines via qPCR. Fibroblasts treated with Snail Tg keratinocytes CM resulted in a significant upregulation of CXCL1, IL6, and IL17F (Figure 3c). It is important to note, however, that the conditioned media was not able to fully recapitulate the inflammatory cytokine profile found in the transgenic skin. It is likely that these cytokines are upregulated by other immune cells, keratinocytes or endothelial cells in the skin, which are implicated in contributing to dermal fibrosis as well but are not represented in the *in vitro* fibroblast culture.

These results beg the question as to the identity of the factor(s) in the Snail Tg conditioned media that are responsible for fibroblast activation. We have previously reported that Fibulin-5 and PAI-1 are secreted by Snail Tg keratinocytes that contribute to fibrosis by altering tissue stiffness (2) and mast cell recruitment (17), respectively. However, we found that ablating these genes from Snail Tg background only results in a partial reduction of the fibrotic phenotype. This incomplete abrogation of the fibrotic phenotype implies that other factors are involved in fibrogenesis each contributing in an additive or synergistic manner. Mindin is a F-spondin family protein which has previously been reported to be overexpressed in some solid tumors (41),(42),(43),(44) and fibrotic conditions (45),(46). Since the Snail tg mice also have features of cutaneous squamous cell carcinoma (40), we hypothesized that Mindin may play a role in Snail mediated fibrogenesis of the skin. Interestingly, we found that Mindin is overexpressed in the dermis in biopsies from SSc patients (Figure 3d). Moreover, Mindin RNA is exclusively expressed by Snail Tg but not WT epidermal keratinocytes (Figure 3e). Since Mindin is known to promote recruitment of macrophages (47),(48), priming of T cells (49), and activation of the pro-inflammatory signaling pathway utilizing NFκB (45),(50) we postulated that Mindin may be required for recruitment of immune cells into the Snail Tg skin and contributed to an inflammatory microenvironment that is crucial for fibrogenesis. In this regard we found that ablating Mindin expression in the Snail Tg background resulted in a drastic reduction in the number of infiltrating T cells and macrophages in P9 skin (Figure 3f). Interestingly, CD3 positive cells which were majorly present in the epidermal compartment of the WT skin were reduced in the epidermis of the Snail Tg skin, which appears to be compensated with an increase in the dermal population of these cells. However, there was no reduction in the number of mast cells in Snail Tg/Mindin KO skin (Supplementary Figure 3d), which we have previously shown is regulated by PAI-1 (17).

As expected with the recruitment of immune cells, inflammatory cytokine expression was also upregulated in the skin of the Snail Tg mouse (Figure 3g). In the absence of Mindin, cytokines in the P9 Snail Tg skin such as IL10, CXCL1, CCL20 were significantly downregulated. While IL6, IL-1β and S100a9 also exhibited a downward trend, it did not reach the statistical significance of p<0.05. The reduction in T cells and macrophages was only apparent in P9 skin. By the age of P60, other inflammatory signals in the Snail Tg skin may override the early protection provided by removing Mindin in the transgenic background (Supplementary Figure 3e). Consistent with this, the absence of Mindin in the skin of P60 Snail Tg mice did not impact the elevated levels of the inflammatory cytokines (Supplementary Figure 3f).

### Mindin stimulates pro-inflammatory cytokines in fibroblasts that drives fibrogenesis

The dependence of cutaneous inflammation on Mindin in the Snail Tg mouse is consistent with Mindin’s known role as a chemoattractant for macrophages (48). We postulated that since Mindin is secreted from transgenic epidermal keratinocytes and diffuses throughout the dermis, it may need a relay system to recruit immune cells from the circulation. One such link between epidermis and vasculature may be the activated fibroblasts which have emerged as an important signaling node for coordinating various processes associated with fibrogenesis (51). In particular myofibroblasts secrete a number of inflammatory cytokines and chemokines. However, it is not known if Mindin can affect fibroblasts and induce them to contribute proinflammatory signals important for the development of fibrosis.

To examine the potential effect of Mindin on fibroblasts, we treated primary dermal fibroblasts with recombinant Mindin and performed an RNA seq analysis. We found 1232 genes were upregulated, and 1274 genes were downregulated (Padj < 0.001, Log2FC>0.5 or Log2FC<-0.5), (Supplementary Figure 4A). In line with the postulated role of Mindin in the recruitment of immune cells, we found that the majority of GO terms enriched for upregulated genes is related to inflammatory processes (Figure 4a). Of these, many cytokines that were upregulated are involved in macrophage, and T cell migration and recruitment (Figure 4b). Thus, while Mindin is known to directly aid in the recruitment of macrophages (48), it may also act on fibroblasts to increase the levels of cytokine expression that can amplify the signal for the migration and homing of immune cells into the skin. In line with this, we found a significant overlap of 16 genes between genes upregulated in Mindin treated fibroblasts and in Snail Tg mouse skin (Supplementary Figure 4a) (representation factor = 1.9, p<0.009). Again, the majority of GO terms that were enriched using the overlapping genes belonged to inflammatory processes (Supplementary Figure 4b). Similarly, there is a significant overlap of 182 upregulated genes between early diffused SSC patients and Mindin treated fibroblasts (Supplementary Figure 4c, d), and an overlap of 44 genes with pediatric localized SSc patients (Supplementary Figure 4e, f). Many genes that we found to be upregulated in the Snail Tg skin via transcriptome profiling were confirmed by qPCR analysis of primary mouse dermal fibroblasts treated with Mindin, such as CXCL1, IL-6, CCL12, CCL-20, IL-1β (Figure 4c).

**Figure 4.**
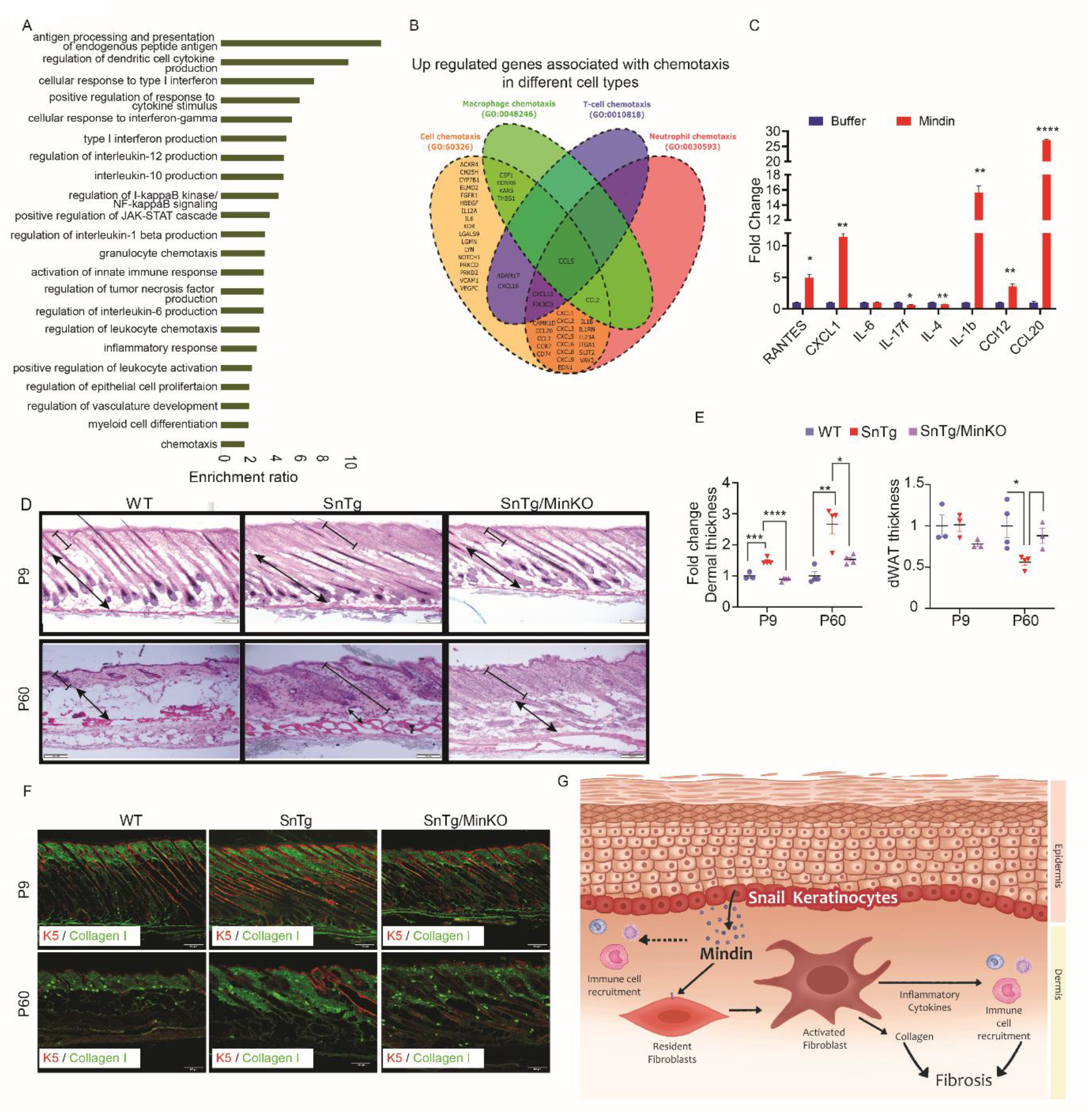
Mindin mediates pro-inflammatory activation of fibroblasts aids in fibrogenesis. A) Biological processes based on GO term enrichment of genes upregulated in fibroblasts treated with Mindin. B) Venn diagram of upregulated genes in Mindin treated fibroblasts that are involved in chemotaxis of various immune cells. C) qPCR for inflammatory cytokine expression in fibroblasts treated with either buffer control (n=3) or Mindin (n=3). D) H & E staining for WT, Snail Tg and Snail Tg/MinKO skin sections taken at P9 or P60. Blocked lines denote dermal thickness and lines with arrowheads denote dWAT thickness. WT and Snail Tg at P9 and P60 are the same samples that were used earlier in Figure 2a E) Quantification of H & E stained section for dermal thickness and dWAT in WT (n ≥3), Snail Tg (n=4) and Snail Tg/MinKO (n=4) mice. WT and Snail Tg at P9 and P60 are the same samples that were used earlier in Figure 2b F) Immunofluorescence for K5 (red) and collagen (green) in WT, Snail Tg and Snail Tg/MinKO skin at P9 and P60. WT and Snail Tg at P9 and P60 are the same samples that were used earlier in Figure 2c G) Model of Snail mediated fibrosis with Mindin involved in fibroblast activation. The error bars represent mean±SEM. P-value was calculated using two-tailed Welch’s t-test (C), and One way ANOVA followed Tukey’s post-hoc analysis for multiple group comparison (E) (* p <0.05, ** p <0.01, *** p <0.001, **** p <0.0001)

Given the importance of inflammation in fibrogenesis, we assessed whether dermal fibrosis would also be affected in the Snail Tg mouse in the absence of Mindin (Figure 4d). In neonatal skin, we observed that the three-fold increase in dermal thickness in the Snail Tg skin was reduced to wild-type levels upon ablation of Mindin (Figure 4d, e). Even in adult skin (P60), the 3.2-fold increase in dermal thickness in the Snail Tg skin was substantially reduced, though it remained slightly higher than wild type levels. Remarkably, the loss of dermal white adipose tissue seen in the adult transgenic skin is reversed upon deletion of Mindin in the Snail background. Consistent with these observations, levels of dermal collagen were reduced upon deletion of Mindin in the Snail background in both neonatal and adult animals (Figure 4f). In addition, as shown in supplementary Table 1, many of the symptoms associated with scleroderma that are recapitulated in the Snail Tg mouse, are substantially reduced when Mindin is removed from the transgenic background. Altogether these data reveal that Mindin plays a critical role in Snail mediated fibrogenesis, as absence of this matricellular protein largely attenuates the fibrotic phenotype.

## DISCUSSION

We report a mouse model based on the transgenic overexpression of Snail in the basal epidermal keratinocytes to mimic its expression in the SSc skin (Figure 4h). The K14-Snail Tg mouse recapitulates the early manifestation and progressive development of a SSc like disease and builds upon our previous work documenting dermal fibrosis in this mouse model (2) (17). Though SSc often develops in the digital extremities and has become a benchmark for diagnosis (8),(52), these criteria have often been overlooked when characterizing animal models of this disease. Interestingly, in line with the symptomatic chronology of SSc, we observed that Snail Tg mice develop swelling of paws and tail as early as postnatal day 5 (P5), becoming most evident by P9. This edematous stage of SSc is often attributed to early inflammatory infiltrates and microvascular defects (53),(54),(55). Concurrent with edema, the vasculature at this stage is also known to undergo spontaneous vasospasms which decrease the blood flow to extremities in response to cold or emotional stress manifested as reversible discoloration of the tissue. This discoloration in response to cold (known as Raynaud’s phenomenon) was also observed in Snail Tg mice. Though both paws and tail of Snail Tg mice manifested a Raynaud’s like effect, it was more predominant in the tail. The progressive vasculopathy which induces the narrowing of blood vessels and the underlying fibrosis that develop in SSc patients often leads to ischemic lesions and necrotic ulcers, which if not attended to, may lead to autoamputation and loss of parts of the digits. While necrotic lesions and autoamputation were quite common in the tail of Snail Tg mice, loss of digits in the paws was not observed. The difference between the animal model of SSc and the human disease might be one of the species specific anatomical differences where the same symptom manifests slightly different in each species.

The early stage of pulmonary involvement in SSc is characterized by microvascular damage in lungs which may contribute to immune infiltration. While we documented increased vascular leakage in lungs of Snail Tg animals, we have not observed progression to pulmonary fibrosis. It is possible that they need more time for the development of fibrosis. Another likely possibility is that other genetic or environmental components are also required for development of pulmonary fibrosis, which may also explain why all patients with SSc do not develop lung fibrosis (56) (57). Another common lung complication in SSc is pulmonary arterial hypertension (PAH). However, due to technical limitations we were not able to probe for development of PAH in the Snail Tg animals and this would be an avenue to pursue in future studies. Nevertheless, one year old Snail Tg animals did exhibit cardiac involvement marked by enlarged size, increase in left ventricle wall thickness, and decrease in volume at systole. There was a significant increase in ejection fraction and fractional shortening which is usually observed in patients with hypertrophic cardiomyopathy, one of the known cardiac abnormalities observed in SSc patients. Interestingly, we previously found that the Snail Tg skin primed the organ for tumor formation (40), which is consistent with reports indicating that SSc patients are more prone to the development of skin cancer (58).

Of the seven criteria of ACR EULAR 2013, the Snail Tg mouse matched at least five - puffy fingers, Raynaud’s phenomenon, necrotic tip ulcers, presence of anti-nuclear antibodies and thickening of the skin especially in the paws and tail. While the leather like appearance (lesion) of the skin on the paws and tail was apparent, it is difficult to clearly score sclerodactyly due to the natural curling of the murine paws. Similarly, red spots and sores were observed in the paws and limbs of Snail Tg mice and in later stages on lesions around the dorsal neck area. However, whether they represent telangiectasia will require further evaluation. Moreover, the thickness of mouse nails makes it challenging to observe nail fold capillaries and hence was not used as an evaluating parameter for the benchmarking to the human disease.

Given the striking similarities between the Snail Tg mouse and human SSc, we utilized this mouse model to further decipher the molecular mechanisms underlying SSc. It is noteworthy that while expression of Snail RNA levels was consistently higher in Snail Tg mice, Snail protein expression was only observed at the early neonatal age (P9). This is likely attributable to the inherent instability of the Snail protein (18). Importantly, this indicates that even transient expression of this transgene was sufficient to set in motion a self-sustaining program that would ultimately manifest as an SSc-like disease. Investigation of the early events that stimulate this self-sustaining program via transcriptome profiling revealed a substantial amount of genes that are shared between the mouse model and the SSc skin and are involved in the activation of both the innate and adaptive immune systems.

The robust inflammatory phenotype in the Snail Tg mouse begs the question of how the expression of Snail in epidermal keratinocytes leads to the dermal infiltration of immune cells. Owing to its canonical role of inducing an epithelial to mesenchymal transition (EMT), a straightforward hypothesis would be that Snail induced an EMT of epidermal keratinocytes into myofibroblasts capable of secreting inflammatory cytokines. Surprisingly, lineage tracing studies clearly demonstrated a non-EMT function for Snail in cutaneous fibrosis. This is consistent with reports of an EMT-independent function for Snail in both lung fibrosis (59) and in pancreatic cancer (60). Instead, we found that Snail initiates an epithelial-mesenchymal crosstalk (Figure 4h). A central player in mediating this intercellular interaction is the protein Mindin. Though it is well-known as a matricellular protein (48) it can also participate in cell signaling pathways such as a pathogen associated molecular pattern (PAMP) to directly recruit macrophages (61). Herein, we present evidence that Mindin can activate primary dermal fibroblasts leading to increased collagen secretion and the expression of inflammatory cytokines and chemokines that can recruit both adaptive and innate immune cells to the skin. The latter observation is consistent with a previous finding that Mindin can activate pro-inflammatory signaling through the NFκB pathway in HK-2 cells (42). Furthermore, we found that Mindin is essential for the early inflammatory response in the Snail Tg mouse. Crippling this cutaneous inflammation via the genetic knockout of Mindin completely abrogated the dermal fibrosis in the neonatal (P9) Snail Tg mouse. That said, the temporal analysis of the Snail Tg phenotype has revealed an intriguing aspect of fibrosis. In contrast to the neonatal Snail Tg mouse, as the animal aged to adulthood, the absence of Mindin in the P60 mouse did not sustain the reduction of cutaneous inflammation. If anything, it was slightly higher in the Snail Tg/Mindin KO mouse compared to the Snail Tg mouse. Nevertheless, there was still a substantial decrease in dermal thickness in the absence of Mindin in the adult Snail Tg mouse, albeit not to levels comparable to wild type skin. Similarly, the loss of dermal white adipose tissue (dWAT) in the adult transgenic mouse was significantly reduced in the absence of Mindin. Together, these observations suggest that inflammation is more important at the initial stages of fibrogenesis rather than the later stages of the pathogenesis or in the maintenance of this pathology.

We have previously reported an important role for fibulin-5 (2) and PAI-1 (17) in dermal fibrogenesis in the Snail Tg skin. However, the effect of Mindin in abrogating the fibrotic phenotype in the Snail Tg mouse is far more pronounced than removing either fibulin-5 or PAI-1. Thus, this data suggests that specific targeting of Snail and/or Mindin has the potential to fill the gap in effective therapeutic targets to halt the progression of SSc. A note of caution is worthwhile as all fibrotic tissues may not be completely the same. For instance, Mindin has been implicated to be anti-fibrotic in liver steatosis (62) and possibly in cardiac hypertrophy (63).

## Material and methods

### Animal studies

C57Bl6 (WT) mice were obtained from The Jackson Laboratory (Bar Harbor, Maine). Mindin KO mice was obtained from You-Wen He (Department of Immunology, Duke University Medical University Medical Center). The K14-Snail Tg mouse was engineered as described earlier (16). The K14-Snail Tg/Mindin KO mouse was developed by breeding the K14-Snail Tg and Mindin KO mice. All mice were bred and maintained under specific pathogen–free conditions at the BLiSC Animal Care and Resource Centre. Mice were sacrificed at P9 (neonatal) and P60(adult). From these animal skin samples were collected for RNA, protein and OCT or paraffin embedding as required. All animal work was approved by the Institutional Animal Ethics Committee in the CJ lab (INS-IAE-2019/06[R1]). Experiments on mice followed the norms specified by the Committee for the Purpose of Control and Supervision of Experiments on Animals (Government of India). All experimental work was approved by the Institutional Biosafety Committee of inStem.

### Gene expression

Total RNA was extracted from skin or cell lysates using TRIzol Reagent (RNAiso Plus, TaKaRa). From isolated RNA, cDNA was synthesized using PrimeScript kit (Takara) this was followed by quantitative qPCR using Power SYBR Mix (Thermo Fisher Scientific) in a QuantStudio™ 5 Real-Time PCR System, 384-well (Thermo Fisher Scientific). *GAPDH* and *β-actin* were used as reference for normalization. The primer sequences used are listed in Supplementary Table 2.

### Human systemic scleroderma skin

Skin-punch biopsies were taken from the arms of patients diagnosed with diffuse systemic sclerosis or non-systemic sclerosis. RNA was isolated from skin samples and subjected to *SNAIL* and *MINDIN* gene expression qPCR analysis. The characteristics of the patients’ samples are provided in Supplementary Table 3.

### Transcriptome data analysis

The differential gene expression analysis between the control and Snail Tg mouse tissue samples were performed by using the CLC genomics workbench differential expression analysis tool (64). The functional and pathway enrichments of the gene ontology terms and KEGG pathways (65) associated with the differentially expressed transcripts was analyzed through WebGestalt (WEB-based GEne SeT AnaLysis Toolkit) tool (27). The p-values were corrected by Benjamini and Hochberg FDR (false discovery rate) correction method and the FDR< 0.05 was considered as statistically significant (66). The overlapping genes were discovered and venn diagrams were obtained by using Venny 2.1 (67). Representation factor and p value for gene overlap was calculated using exact hypergeometric probability using the program by Jim Lund available at (http://nemates.org/MA/progs/representation.stats.html).

### Western blot

The tissue samples were crushed using mortar and pestle, samples were prepared in RIPA buffer with protease inhibitors (Sigma, #P2714) these samples were sonicated at 4°C. Tissue lysate was quantified for protein using GeNei BCA protein estimation kit. Using a 4X Laemmli sample buffer, protein samples were prepared and heated at 95°C for 5 min before loading on the polyacrylamide gel. This was later transferred onto a nitrocellulose membrane. The following primary antibodies were used at a dilution of 1:1,000: HA antibody (Abcam ab 9110); 1:4,000 b-actin antibody (Sigma A2228). The HRP-labeled secondary antibodies (Jackson ImmunoResearch) were used at 1:5,000 dilution. Blots were developed on an ImageQuant LAS4000.

### Immunostaining and histology

Skin tissue pieces were either fixed in Bouin’s solution, dehydrated, and embedded in paraffin or embedded directly in tissue freezing medium (Leica). For staining the OCT sections, 10μm thickness were fixed in 4% paraformaldehyde (PFA) followed by blocking before proceeding to the primary antibody staining. Following primary antibodies were used at a dilution of 1:200: K5 (generated in-house); 1:200: Collagen1 (Abcam ab21286); 1:200: CD11b (Abcam ab8878); 1:200: CD3 (e-biosciences 14-0041-85); 1:200: Fibronectin (Abcam ab2413). Alexa Fluor 488– or Alexa Fluor 568–labeled secondary antibodies (Jackson ImmunoResearch) were used at a dilution of 1:300. Hoechst stain was used to mark nuclei. For secreted protein staining, the use of detergent was completely avoided, and K5 staining was used as an internal control. For IHC 3% H_2_O_2_ incubation was done after primary antibody, and HRP-labeled secondary antibodies (Jackson ImmunoResearch) was used. Development was done using DAB substrate (Vector Laboratories; SK4105).

For H&E staining, sections were thawed at room temperature. Sections were stained with hematoxylin for 15-20 seconds, excess stain was removed by dipping the slides in tap water, then eosin was added for 5 seconds and excess stain was removed by dipping the slides in distilled water. Dermal thickness was quantified from the average of multiple measurements made along the length of the H&E-stained sections. For Oil red O staining fix the slides with 4% PFA wash with PBS. Immerse the slides in freshly prepared 60% isopropanol for 2-3 seconds. Add oil red O for 60-70 seconds. Immerse the slides in 60% isopropanol for 5 seconds. Dip the slides in water to wash off excess stain.

Toluidine blue staining was performed with 1% toluidine blue solution in 70% ethanol, diluting the solution to 1:10 in a 1% sodium chloride solution, pH 2, for 5 to 30 minutes, followed by extensive washing with water.

b-Gal staining was performed by fixing OCT sections using 0.5% glutaraldehyde solution for 2 mins at room temperature. Washed with 0.1 M sorensen’s phosphate buffer (0.02M sodium phosphate monobasic and 0.08M sodium phosphate dibasic). Incubate skin sections with X-Gal staining solution at 37°C overnight. Terminate by incubating in sorensen’s phosphate buffer.

Imaging was done on an Olympus IX73 microscope or FV1000 confocal microscope and analyzed on the Fiji software.

### Raynaud’s assay

P9 (neonatal) mice tails were kept on ice for approximately 2 minutes after which discoloration was observed in Snail Tg mice as compared to WT. Images were captured with a camera immediately after the cold exposure and again after 24 hours.

### Anti-nuclear antibody staining

Serum from WT and Snail Tg mice was collected from the blood. This serum was added on fixed cultured primary mouse fibroblasts followed by permeabilization and incubated for 2 hours at room temperature. Anti-mouse secondary antibody (Jackson ImmunoResearch) was used to observe nuclear localization

### Vascular leakage analysis

Evans Blue dye (Sigma, E2129) (1% w/v in PBS) was injected retro-orbitally in wildtype and Snail Tg mice according to 60mg/kg body weight of the animals. Lungs were harvested after euthanizing the animals 30 mins post-injection. Lung tissues were divided into 3 tubes for each animal. 500µl formamide was added to each tube after air-drying. Tubes were incubated in a 55 °C water bath for 48 hours to extract dye from the tissue. Absorbance was measured at 620nm using formamide as blank. Ng of Evan’s Blue extravasated per mg tissue was calculated from a standard curve.

### Echocardiography imaging and analysis

To assess cardiac function in WT and Snail Tg mice at one year of age, non-invasive M-mode echocardiography was performed using a VisualSonics Vevo 3100 imaging system (FUJIFILM VisualSonics Inc.) with an MX400 20–46 MHz transducer. Under isoflurane anaesthesia delivered via inhalation (3% for knockdown and 0.5 to 1% for sedation during the duration of the procedure), the hearts were monitored to assess function and morphology. Two-dimensional images and M-mode tracings were recorded on the parasternal long axis at the level of the papillary muscle to determine the percentage EF, FS and ventricular dimensions. Data were analysed using Vevo LAB version 3.1.0 (Build 13029).

### Lineage tracing experiment

R26R mice were crossed with K14-Cre mice to generate K14-Cre R26R mice. K14-Cre R26R were crossed with K14-Snail Tg mice to generate K14-Cre R26R/K14-Snail Tg mice. Skin sections were taken from K14-Cre R26R and K14-Cre R26R K14-Snail Tg mice and probed for expression of β-Gal and αSMA using IHC.

### Cell Culture

Primary dermal fibroblasts were isolated and cultured as described earlier (2). Fibroblasts were cultured in DMEM high glucose media with 10% FBS.

Primary epidermal keratinocytes from WT and Snail Tg mice were isolated and cultured as described earlier (68). The keratinocytes were grown in low calcium (50uM) E-media, and after reaching confluency cells were washed with PBS. Cells were then incubated with P-media (E-media without serum) for 1 hour and the media was removed. P-media was again added on WT and Snail Tg keratinocytes and incubated for 16 hours to generate the conditioned media.

Primary fibroblasts were treated with conditioned media from WT or Snail Tg keratinocytes or Mindin for 16 hours. RNA was then extracted from these cells using TRIzol Reagent.

Mindin expressing CHO cells were provided by You-Wen He (Department of Immunology, Duke University Medical University Medical Center). These cells were cultured in DMEM high glucose media with 10% FBS.

### Mindin purification

Histidine-tagged Mindin was purified from conditioned media collected from CHO-Mindin cells using Ni-NTA beads.

### Collagen contraction

Rat tail collagen plugs (MilliporeSigma; 08-115; 1 mg/ml) were made in 5% FBS-containing media with 100,000 fibroblasts. Each plug was kept in 500 μl media with buffer or conditioned media. Contraction was quantified by calculating the area of the gel after 72 hours.

### Statistics

Comparisons of 2 groups were done using a 2-tailed, Welch’s t test or Mann Whitney U-test. One-way ANOVA followed by Tukey’s post hoc analysis was used for multiple group comparisons. GraphPad Prism 6 (GraphPad Software) was used for all statistical analyses.

Data represents the mean ± SEM. P values of less than 0.05 were considered significant.

### Study approval

Animal work conducted at the NCBS/inStem Animal Care and Resource Centre was approved by the inStem Institutional Animal Ethics Committee following the norms specified by the Committee for the Purpose of Control and Supervision of Experiments on Animals (Government of India). Acquisition and processing of the human tissue were conducted according to the protocol approved by the IRB of the CMC. Informed consent was obtained from all patients for skin sample collection and experimentation. All experimental work was approved by the Institutional Biosafety Committee and the Institutional Human Ethics Committee of inStem.

## Acknowledgements

The authors would like to thank members of Jamora laboratory for their critical review of the work and insightful discussions, Binita Dam, Abrar Rizvi, Syed Shahid Musvi, Yogesh Chandra for their technical assistance, and Ritoparna Hazra for designing the graphical model. This work was supported by core funds from the Institute for Stem Cell Science and Regenerative Medicine (inStem), and grants from the Department of Biotechnology of the Government of India (BT/PR8738/AGR/36/770/2013) and (BT/PR32539/BRB/10/1814/2019); the National Institute of Arthritis and Musculoskeletal and Skin Diseases (NIAMS), NIH (5R01AR053185-03); and the American Cancer Society (15457-RSG-08-164-01-DDC) to CJ; PSD is supported by the DBT/Wellcome Trust-Indian Alliance (IA/I/16/1/502367). IR and EYH were supported by the funding of the Indian Council of Medical Research (Senior Research Fellowship); SK was supported by the funding of the National Centre of Biological Sciences. Animal studies were partially supported by the National Mouse Research Resource (NaMoR) grant BT/PR5981/MED/31/181/2012;2013-2016;2018 and 102/IFD/SAN/5003/2017-2018 from the Department of Biotechnology. We thank the staff of the BLiSC Animal Care and Resource Centre for assistance with animal husbandry, and the BLiSC Central Imaging and Flow Cytometry Facility for help with image acquisition

## Author contribution

IR, SK, TLT designed and performed experiments, evaluated and interpreted data, and wrote the manuscript. IR, SK, TLT, EYH, DKK, DS, JA, ASHPA, RFZ, HJ performed experiments. SP, PK, RD, SJ performed bioinformatics analysis. RS, RG, DD, and PMJ provided the human scleroderma samples. DP provided guidance in cardiac and bioinformatic analysis. JV provided guidance, and reviewed the manuscript.YWH provided Mindin Ko mice and CHO-Mindin cell line. CJ helped in designing the experiments, provided guidance, and reviewed the manuscript.

### Conflict of interest

The authors have declared that no conflict of interest exists

## Supplementary Figures

**Supplementary Figure 1.**
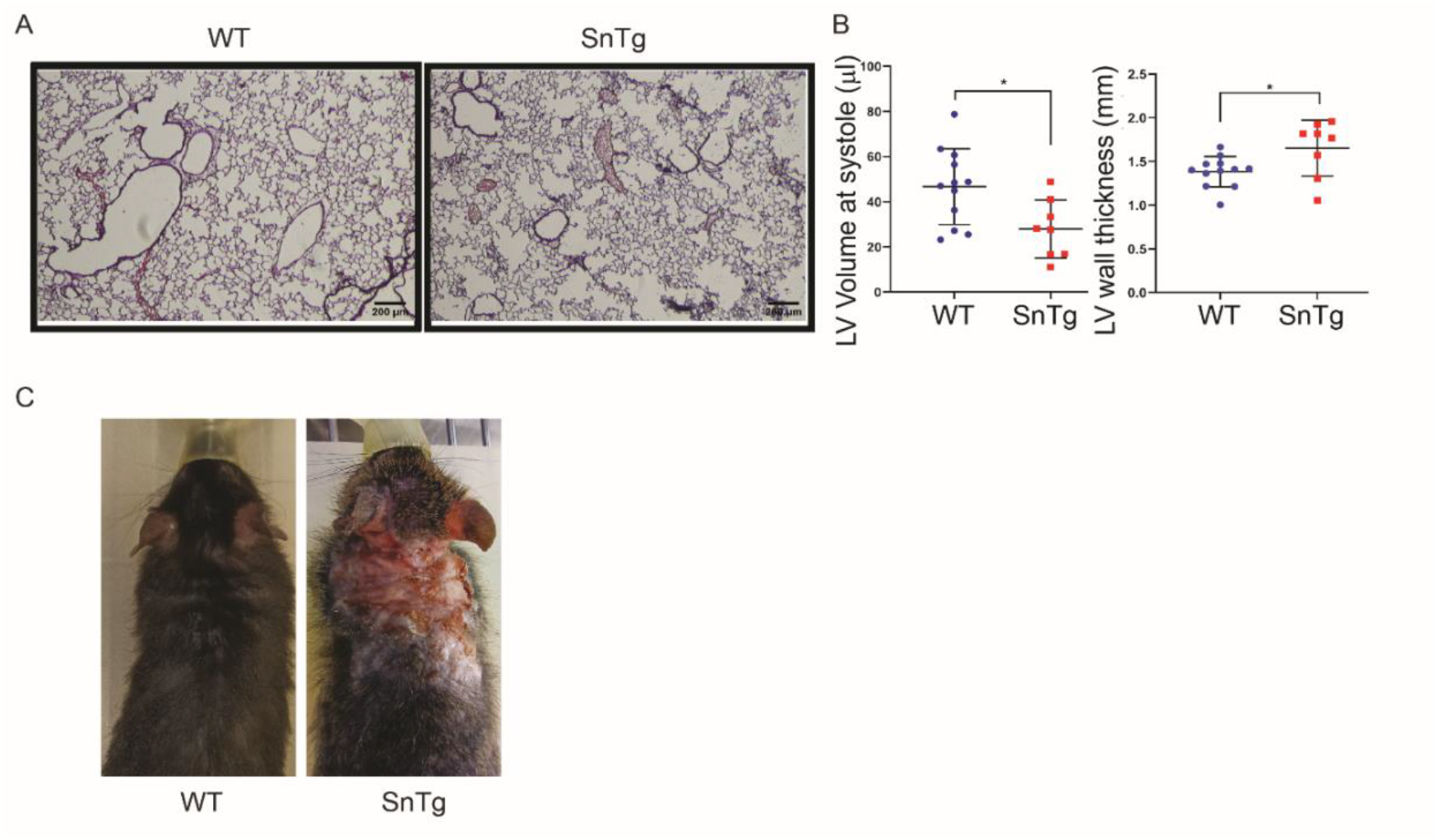
Snail Tg mouse model recapitulates diagnostic feature of scleroderma. A) H & E staining of lung sections taken from WT and Snail Tg mice at P60 B) Echocardiogram of the heart in 1-year old WT and Snail Tg mice. Comparison of values for LV volume at systole and LV wall thickness between WT (n=12) and Snail Tg (n=8) mice. C) Gross appearance of the skin lesions observed at the dorsal neck area of the adult (P90) Snail Tg mice (Right panel) compared to the WT (P90) (Left panel). The error bars represent mean±SEM. P-value was calculated by Welch’s t-test (B). (* p <0.05).

**Supplementary Figure 2.**
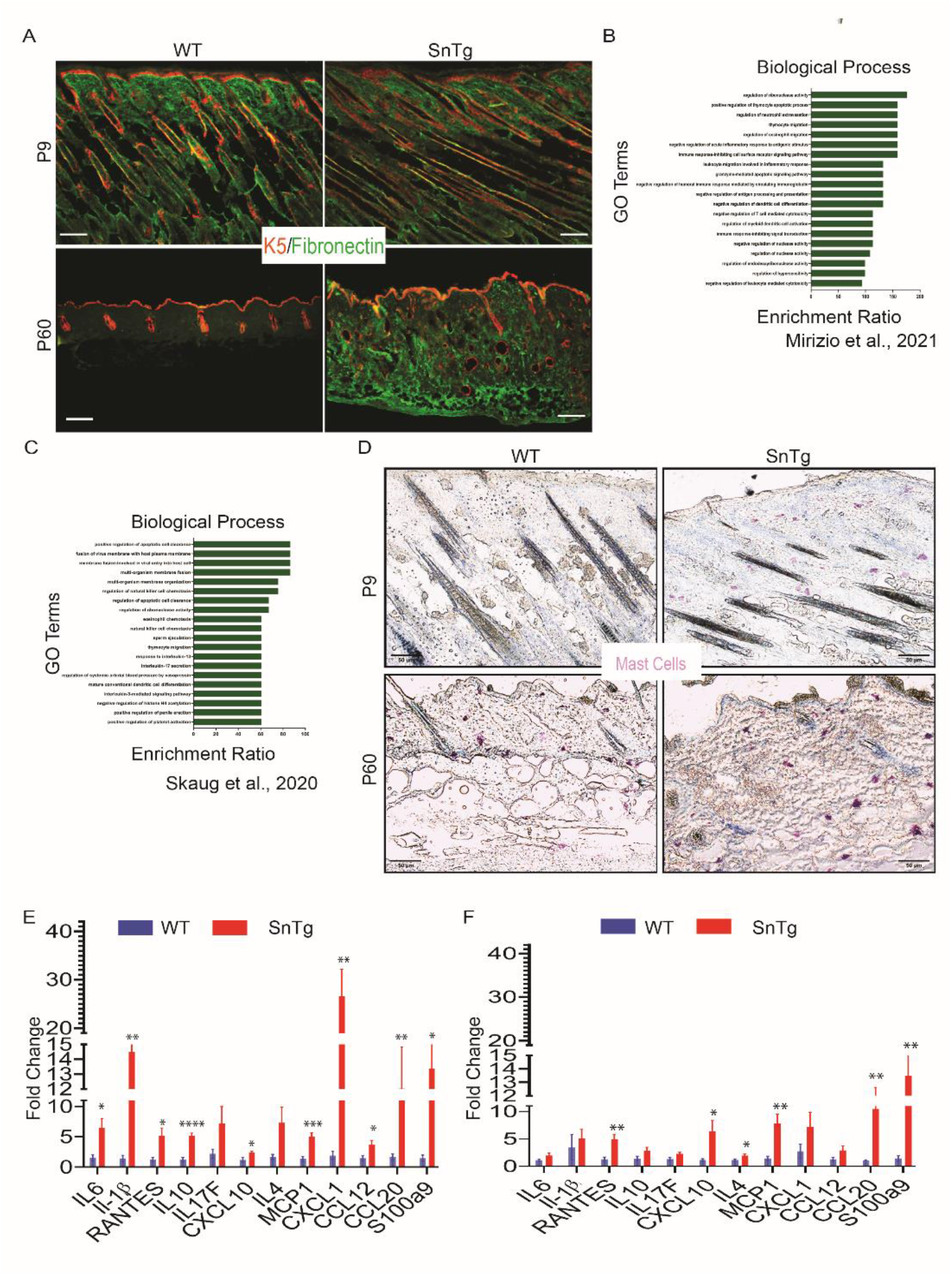
Snail Tg mice recapitulates histological and molecular characteristic of SSc. A) Immunofluorescence imaging for K5 (red) Fibronectin (green) in WT and Snail Tg mice at P9 and P60. B) Biological processes based on GO term enrichment of upregulated genes common in skin of the Snail Tg mice and SSc (GSE166861 (28)). C) Biological processes based on GO term enrichment of upregulated genes common in skin of the Snail Tg mice and SSc (GSE130955, (27)). D) Toluidine blue staining for mast cells in WT and Snail Tg at P9 and P60 E) qPCR for inflammatory cytokines gene expression in WT (n=8), Snail Tg (n≥6) skin at P9. F) qPCR for inflammatory cytokines gene expression in WT (n=9), Snail Tg (n=7) skin at P60. The error bars represent mean±SEM. P-value was calculated using two-tailed Welch’s t-test (E,F). (* p <0.05, ** p <0.01, *** p <0.001, **** p <0.0001).

**Supplementary Figure 3.**
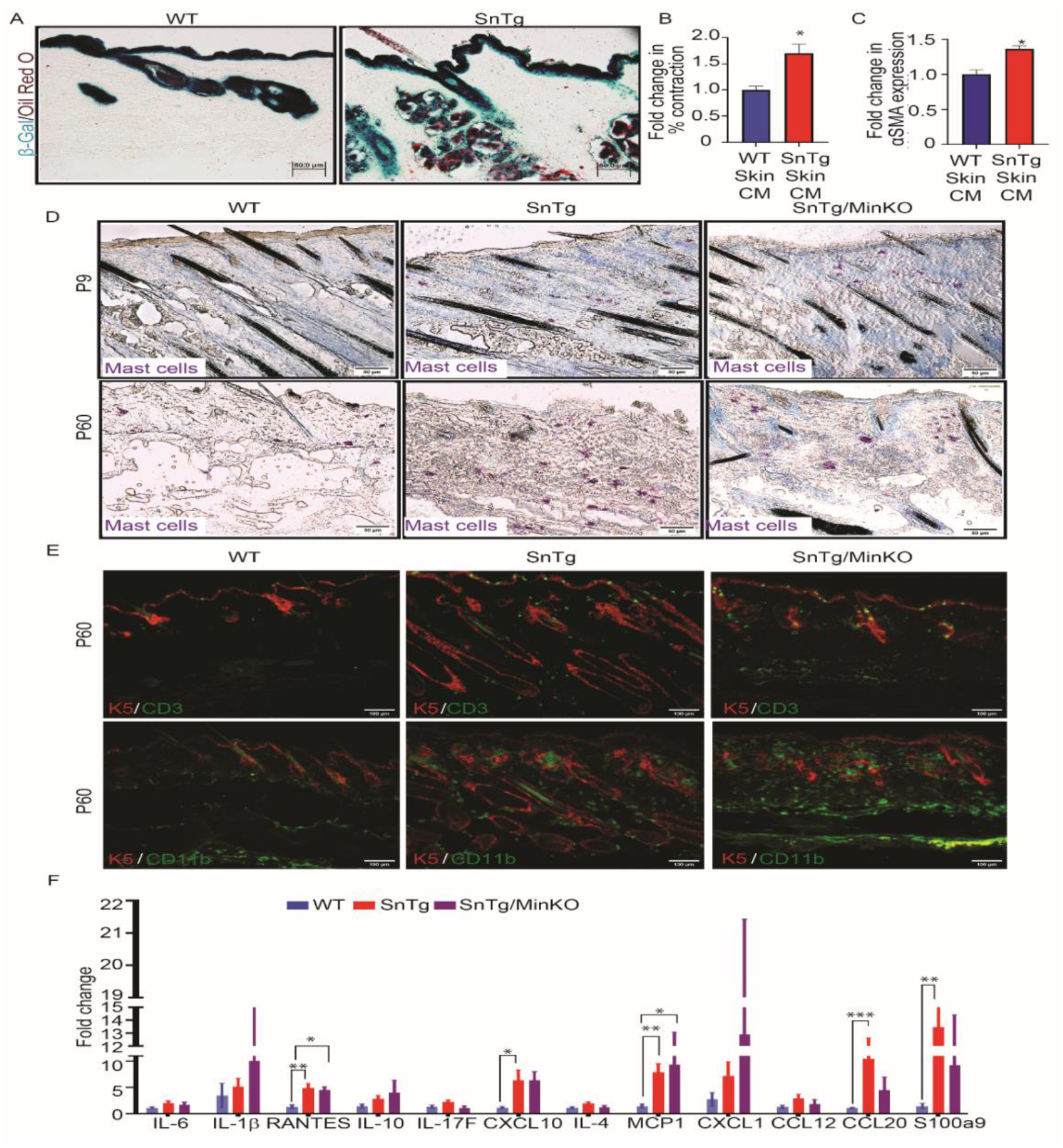
Secreted Mindin leads to fibrotic phenotype. A) Lineage tracing of the WT and Snail Tg mice using the K14-Cre R26R reporter. All cells of epithelial origin are positive for β-Gal (blue) and oil red O marks the sebaceous glands (red). B) Conditioned media (CM) was collected from WT (n=3) and Snail Tg (n=3) mouse skin explants. Fibroblast activation measured with the collagen contraction assay. C) αSMA gene expression to assess fibroblasts activation after treatment with WT and Snail Tg skin conditioned media D) Mast cells staining in WT, Snail Tg and Snail Tg/Mindin KO at P9 and P60. WT and Snail Tg for P9 and P60 are the same samples used earlier for Supplementary Figure 2d E) Immune cell populations in WT, Snail Tg, and Snail Tg/MinKO mice skin at age P60. Skin sections were stained for keratin-5 (K5, red) and either macrophages (CD11b, green) or T-cells (CD3, green). WT(P60) and Snail Tg(P60) are the same samples used earlier for Figure 2h-i F) qPCR for inflammatory cytokines gene expression in WT (n=9), Snail Tg (n=7) and Snail Tg/MinKO (n=3) skin at P60. WT(P60) and Snail Tg(P60) are the same samples used earlier for Supplementary Figure 2f The error bars represent mean±SEM. P-value was calculated using two-tailed Welch’s t-test (B,C), and One way ANOVA followed Tukey’s post-hoc analysis for multiple group comparison (F) (* p <0.05, ** p <0.01, *** p <0.001, **** p <0.0001)

**Supplementary Figure 4.**
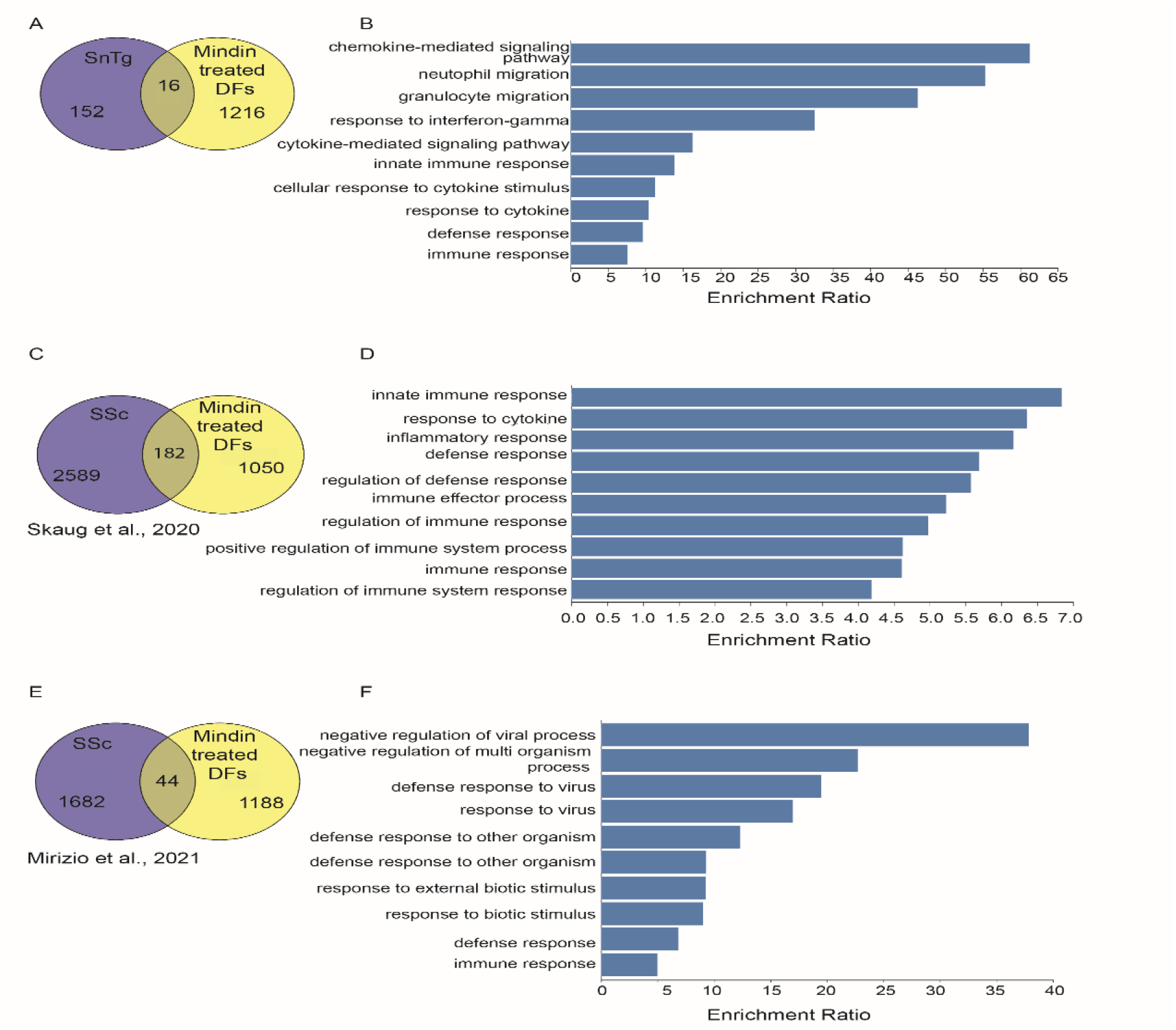
Mindin treated fibroblasts show an overlap with Snail Tg mice and human SSc. A) Venn Diagram of overlapping upregulated genes in Snail Tg mice and Mindin treated fibroblasts. B) Top 10 biological processes GOTERMS of upregulated genes that are held in common in Snail Tg mice and Mindin treated fibroblasts. C) Venn Diagram of overlapping upregulated genes in early dSSc patients [GSE130955, (27)] and Mindin treated fibroblasts. D) Top 10 biological processes GOTERMS of upregulated genes that are held in common in dSSc and Mindin treated fibroblasts. E) Venn Diagram of overlapping upregulated genes in early pediatric Limited SSc patients [GSE166861 (28)] and Mindin treated fibroblasts. F) Top 10 biological processes GOTERMS of upregulated genes that are held in common in pediatric Limited SSc and Mindin treated fibroblasts.

**Supplementary Table 1:** Number of animals exhibiting the noted symptom/total number of animals observed.

| **P9** | sample size | Raynaud's phenomenon in tail | Puffy paws/tail | Skin lesions on tail | Tail tip ischemic ulcer |
| --- | --- | --- | --- | --- | --- |
| WT | 17 | 0/17 | 0/17 | 0/17 | 0/17 |
| Snail Tg | 6 | 5/6 | 4/6 | 5/6 | 3/6 |
| Snail Tg/MinKO | 5 | 2/5 | 1/5 | 4/5 | 2/5 |

| **P60** | sample size | Raynaud's phenomenon in tail | ANA | Puffy paws | Tail autoamputation |
| --- | --- | --- | --- | --- | --- |
| WT | 8 | 0/8 | 0/6 | 0/8 | 0/8 |
| Snail Tg | 9 | 8/9 | 4/7 | 4/9 | 9/9 |
| Snail Tg/MinKO | 5 | 2/5 | 1/4 | 1/5 | 2/5 |

**Supplementary Table 2:**
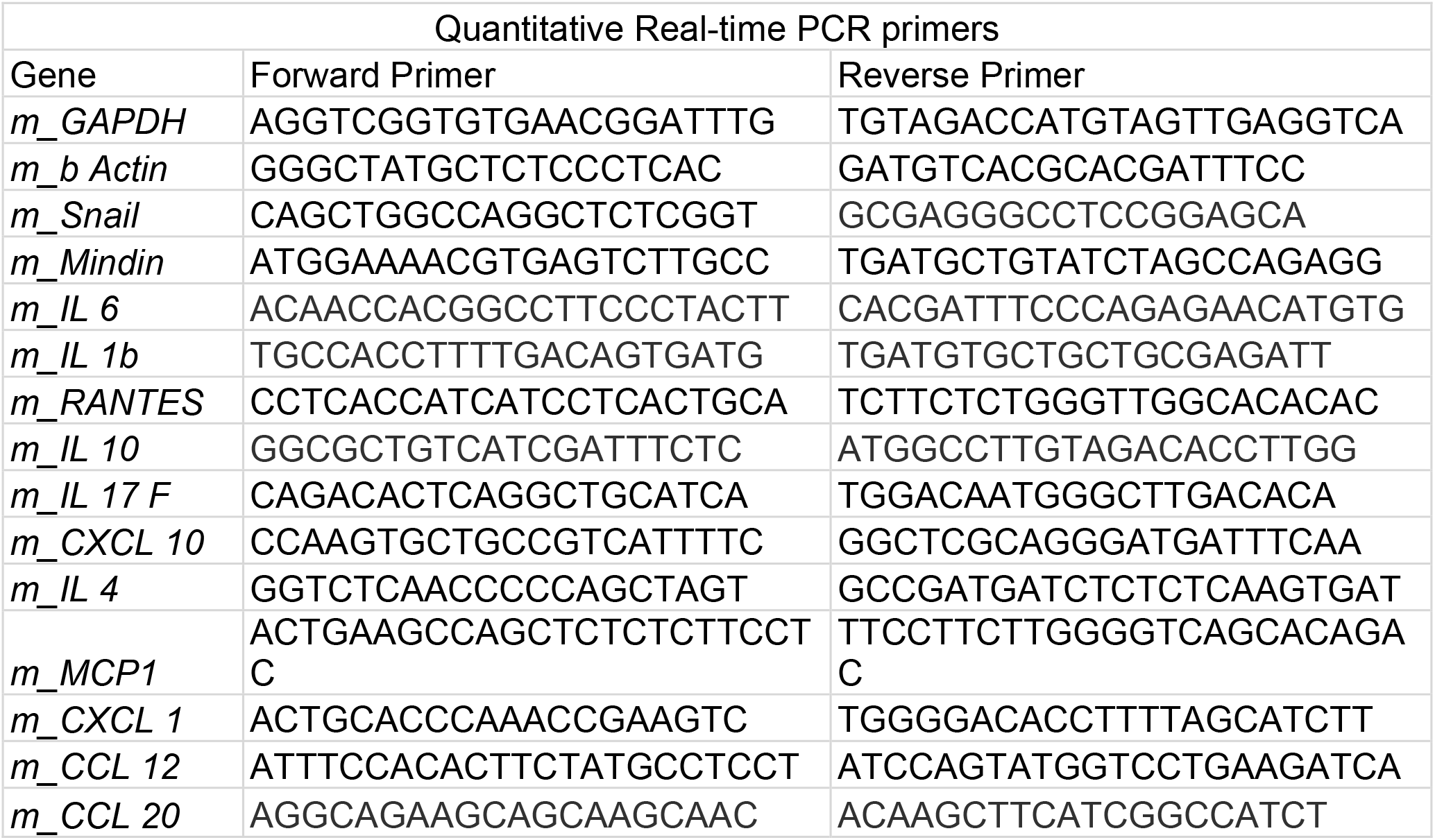
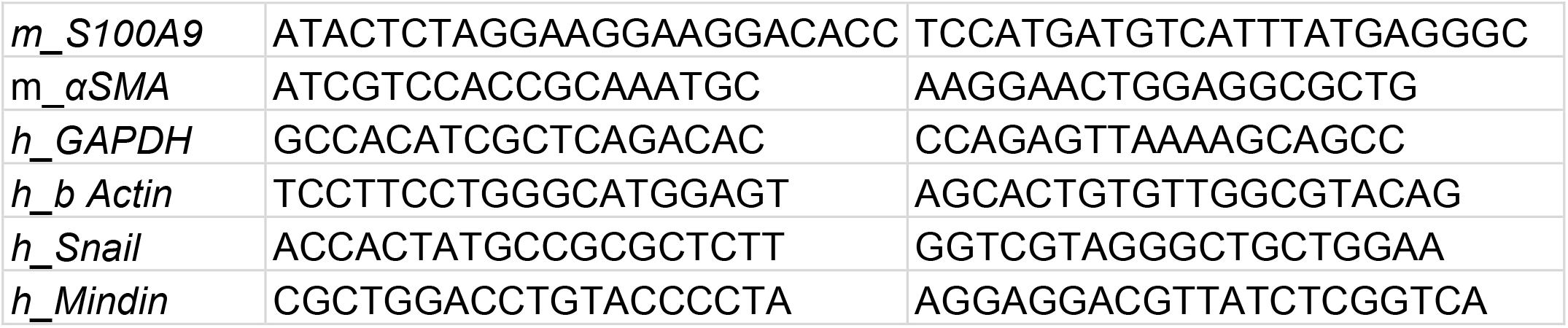
List of primers used for qPCR

**Supplementary Table 3:**
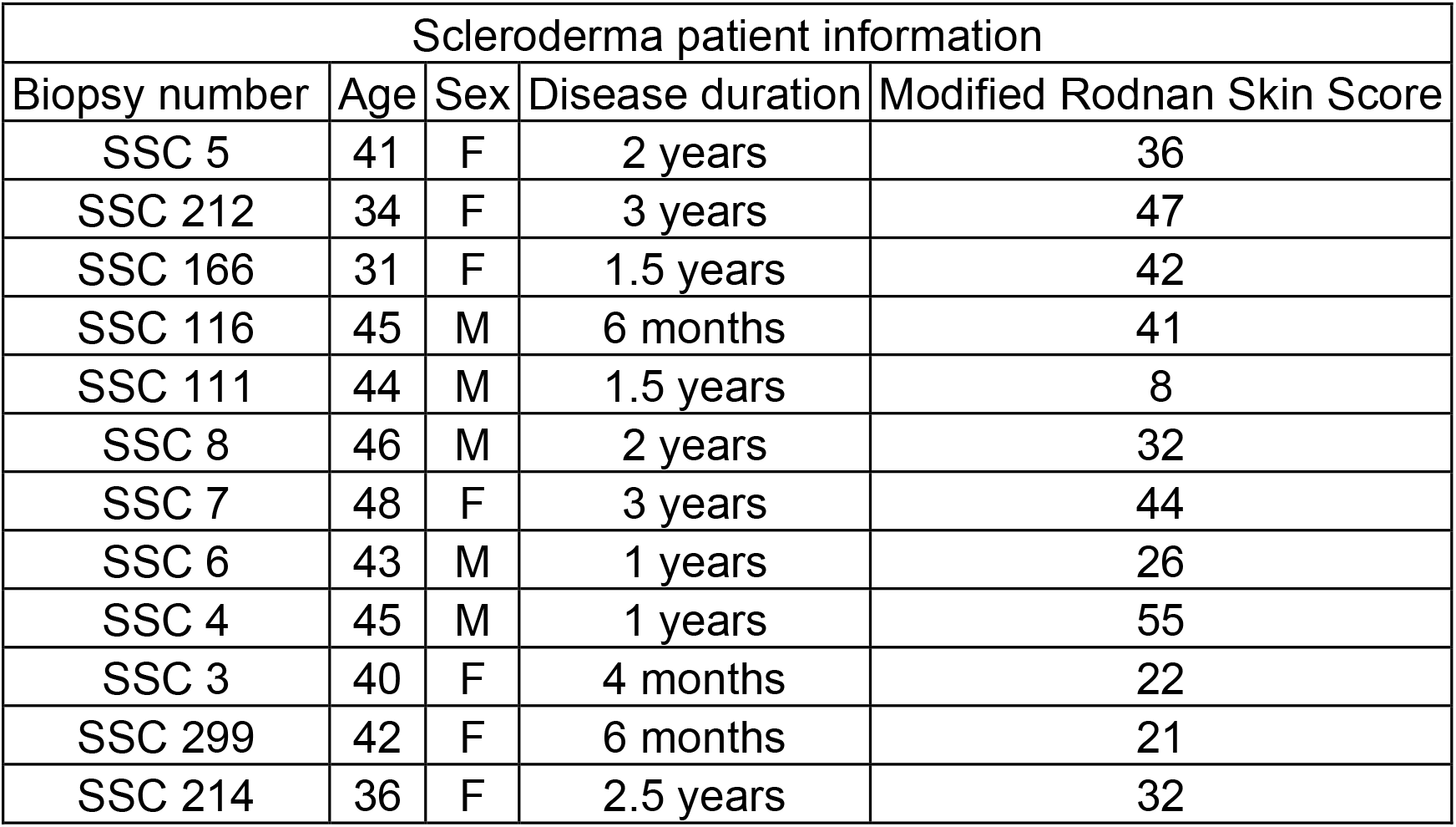
Scleroderma patient information. Non-SSC samples were collected from healthy humans with same age and sex. These samples were the same used in Nakasaki et al., 2015 (2).

| Scleroderma patient information |  |  |  |  |
| --- | --- | --- | --- | --- |
| Biopsy number | Age | Sex | Disease duration | Modified Rodnan Skin Score |
| SSC 5 | 41 | F | 2 years | 36 |
| SSC 212 | 34 | F | 3 years | 47 |
| SSC 166 | 31 | F | 1.5 years | 42 |
| SSC 116 | 45 | M | 6 months | 41 |
| SSC 111 | 44 | M | 1.5 years | 8 |
| SSC 8 | 46 | M | 2 years | 32 |
| SSC 7 | 48 | F | 3 years | 44 |
| SSC 6 | 43 | M | 1 years | 26 |
| SSC 4 | 45 | M | 1 years | 55 |
| SSC 3 | 40 | F | 4 months | 22 |
| SSC 299 | 42 | F | 6 months | 21 |
| SSC 214 | 36 | F | 2.5 years | 32 |

